# Cytogenetic events in the endosperm of amphiploid *Avena maroccana* × *A. longiglumis*

**DOI:** 10.1101/2021.02.10.430611

**Authors:** Paulina Tomaszewska, Romuald Kosina

**Author notes:** Correspondence: R. Kosina, Institute of Environmental Biology, University of Wrocław, Przybyszewskiego 63, 51-148 Wroclaw, Poland.

## Abstract

This study analysed cytogenetic events occurring in the syncytial endosperm of the *Avena maroccana* Gand. × *Avena longiglumis* Dur. amphiploid, which is a product of two wild species having different genomes. Selection through the elimination of chromosomes and their fragments, including those translocated, decreased the level of ploidy in the endosperm below the expected 3*n*, leading to the modal number close to 2*n*. During intergenomic translocations, fragments of the heterochromatin-rich C-genome were transferred to the A and Al genomes. Terminal and non-reciprocal exchanges dominated, whereas other types of translocations, including microexchanges, were less common. Using two probes and by counterstaining with DAPI, the *A. longiglumis* and the rare exchanges between the A and Al genomes were detected by GISH. The large discontinuity in the probe labelling in the C chromosomes demonstrated inequality in the distribution of repetitive sequences along the chromosome and probable intragenomic rearrangements. In the nucleus, the spatial arrangement of genomes was non-random and showed a sectorial-concentric pattern, which can vary during the cell cycle, especially in the less stable tissue like the hybrid endosperm.

## Introduction

Endosperm, which is the storage tissue of grass caryopsis, plays an important role in human nutrition, and hence, should be thoroughly analysed. Many grass species, including cereals, have evolved through the processes of hybridisation and polyploidisation (Grant 1981). In young hybrids, endosperm development is highly unstable as in Triticale (Kaltsikes 1973; Peña et al. 1982) or oats (Tomaszewska and Kosina 2018) and is correlated with the cytogenetic behaviour of the plant. In addition, the low level of telomerase activity in the endosperm, which was first observed in barley and maize, has been found to affect the stability of telomeres and result in cytogenetic aberrations (Kilian et al. 1998). In the young pistils of *Arabidopsis* Heynh., shortened telomeres were shown to induce the breakage/fusion/bridge (BFB) cycle (Siroky et al 2003). Such cytogenetic behaviour was also observed in human cancer cells presenting of telomere instability (Lo et al. 2002).

Most studies available in the literature on cytogenetic data are related only to root tissues and not endosperm. Thus, for instance, in the roots of *Avena sativa* L., intergenomic translocations, which have been visualised as large exchanges using genomic *in situ* hybridisation (GISH) by Chen and Armstrong (1994), but as sites of point hybridisation by Jellen et al. (1994), proved that even small translocations can change the genome status. The level of cytogenetic disorders can be extremely high, as was observed in a single *Elymus farctus* (Viv.) Runemark ex Melderis plant (Heneen 1963). There is a common agreement that heterochromatin is involved in these changes, and in the species mentioned above, large blocks of terminal heterochromatin have been discovered (Endo and Gill 1984). Furthermore, it has been proved that the sites of C-banding (heterochromatin) and translocation breaks coincide in the A and C genomes of *Avena maroccana* Gand. (Jellen et al. 1994). The C genome is highly heterochromatic (Fominaya et al. 1988b), and such a chromosome structure can contribute to translocation events. The C genome of *A. maroccana* also has heterochromatin dispersed along the chromosomes (Shelukhina et al. 2007), while the A genomes of various diploid oats have less heterochromatin localised mainly at telomeres (Fominaya et al. 1988a; Badaeva et al. 2005). This intergenomic difference in the amount of heterochromatin can significantly affect the dynamics of chromosomal rearrangements under the conditions of hybridity. In addition, during the evolution of AACC tetraploids, both A and C genomes were altered to different degrees, especially the C genome (Drossou et al. 2004).

The parental species of the *A. maroccana* Gand. × *Avena longiglumis* Dur. amphiploid are meiotically compatible in the crossing process exhibiting partial sterility (Loskutov 2001), but the result of their crossover is influenced by the ecotype of *A. longiglumis* (Rajhathy 1971). The genomic status of the amphiploid is determined by the C genome and two different A genomes. In the hybrids formed by crossing *Avena strigosa* Schreb. (AsAs genomes), *Avena eriantha* Durieu (CpCp genomes), and *A. maroccana* (AACC genomes), the meiotic division is characterised by a poor pairing of chromosomes, which can be related to translocation variability rather than the lack of chromosome homology (Leggett 1998; Nikoloudakis and Katsiotis 2015). The last statement is very important due to the role played by small and large translocations in stabilising the reproduction system of young polyploid hybrid forms and their speciation.

The present molecular studies on oat genomes proved that the A genome of *A. maroccana* has to be considered as D genome (Yan et al. 2016). This was corroborated by the similarity between the chromosomes 10A ‘magna’ and 21D ‘sativa’ (Fominaya et al. 2017). The principal coordinates analysis showed that *A. eriantha* and *A. longiglumis* are closer to AACC oat tetraploids than other AA diploids (Yan et al. 2016). Both species were accepted as ancestral during the evolution of AACC tetraploids and AACCDD hexaploids (Chew et al. 2016). The exceptional role played by *A. longiglumis* in the oat evolution was also studied by Fu (2018). The author analysed cp and mt genomes and proved that *A. longiglumis* was a maternal component that created *Avena insularis* Ladiz. (AACC), and subsequently a paternal species with *A. insularis* as a maternal species in the evolution of the AACCDD species. Thus, in the light of the above data, the research of a maroccana-longiglumis hybrid unit and its endosperm can significantly supplement the knowledge about the biology of the cereal developing large caryopses.

GISH analyses have revealed that in the sexual hybrids, different genomes are not randomly distributed in the nucleus (Schwarzacher et al. 1989). This non-random architecture of chromatin is maintained throughout the cell cycle (Leitch et al. 1991). Genome separation has also been observed in somatic hybrids, but it was found that there is a change in the pattern of separation from segmental to radial (Gleba et al. 1987). In the case of grass endosperm, the genomes were found to change the arrangement from sectorial to concentric in nuclei approaching apoptosis (Tomaszewska and Kosina 2013). It has been shown that the architecture of genome domains correlates with genes expression (Bennett 1984; Heslop-Harrison 1990). However, the reference data on nuclear architecture in the endosperm are not common.

The endosperm is a complex object to conduct a cytogenetic analysis compared to other tissues due to less chromatin condensation as shown in *Arabidopsis* (Baroux et al. 2007). For this reason, identifying chromosome rearrangements and other cytogenetic aberrations is very challenging. Thus, the overall purpose of this paper is to enrich the available data on endosperm cytogenetics in cereals. More specifically, it focuses on the identification of genomes in the hybrid endosperm and establishing the relationships between them by determining the level of genome rearrangement, with an emphasis on intergenomic translocations.

## Materials and methods

### Plant material

To perform a cytogenetic analysis of the free-nuclear syncytium of the oat amphiploid *A. maroccana* Gand. × *A. longiglumis* Dur. (Am×Al) and its parental species (taxonomic units) young embryo sacs were sampled from plants, that were cultivated on small plots in the grass collection (Wroclaw, SW Poland), which were maintained by R. Kosina. During the plot experiments, the plants were grown under the same soil-climatic conditions. For each taxonomic unit, 45 embryo sacs were mounted on 15 microscopic slides by pooling three on each. Thus, the study material can be considered as a one-way classification in a completely randomised design. Finally, clear stages of mitosis were selected on microscopic slides as small samples (*n* < 30 or larger) for photographic documentation and quantification.

### Cytogenetic preparation

Mitotic spreads were prepared from the endosperm at the syncytial stage. The endosperm was isolated from young caryopses between 2 and 3 days after pollination, and fixed in ethanol:acetic acid (3:1) solution. The fixed endosperm was washed in an enzyme buffer (10 mM citric acid/sodium citrate) for 15 min, and digested with enzyme mixture containing 0.3% cellulase from *Aspergillus niger*, 0.3% pectolyase from *Aspergillus japonicus*, and 0.3% cytohelicase from *Helix pomatia* in the enzyme buffer for 45 min at 37^°^ C. The digested endosperm was washed again in the enzyme buffer for 15 min and squashed in 45% acetic acid. Coverslips were removed after the slides were frozen with liquid nitrogen. Then the slides were air-dried and used for the *in situ* hybridization procedure.

### Probe preparation

Genomic DNAs were extracted from fresh leaves of *A. eriantha* (NSGC PI 657576) and *Avena nuda* L. (NSGC CIav 9010) using a standard method based on cetyltrimethylammonium bromide (CTAB) (Doyle and Doyle 1990) with minor modifications. The obtained DNAs were used as probes for GISH. The DNA of *A. eriantha* was labelled with tetramethylrhodamine-5-dUTP and that of *A. nuda* with digoxigenin-11-dUTP by nick translation, using a commercially available kit (Roche). After labelling, ethanol precipitation was performed. The DNA pellets thus obtained were washed with 70% ethanol prior to drying and resuspended in water.

### Genomic in situ hybridisation

GISH was carried out as described by Schwarzacher et al. (1989) with minor modifications. Prior to the procedure, the slides were fixed in ethanol:acetic acid (3:1) solution for 10 min, and washed two times in 96% ethanol (10 min each). The slides were air-dried and treated with 0.1 mg/ml of DNase-free RNase A in 2×SSC (0.3 M NaCl, 0.03 M sodium citrate, pH 7) for 1 h at 37^°^ C. Then, the slides were washed with 2×SSC for 15 min and 0.01 M HCl for 5 min at room temperature (RT). After washing, 200 µl of 5 µg/ml pepsin in 0.01 M HCl was applied to each slide, and the slides were incubated for 15 min at 37^°^ C. Next, the slides were washed twice with 2×SSC (5 min each) and incubated in freshly prepared 4% paraformaldehyde for 10 min at RT. The slides were again washed twice with 2×SSC (5 min each), dehydrated through a series of ethanol solutions (70%, 85%, 96%; 10 min each) and finally air-dried.

A hybridisation mixture consisting of 50% deionised formamide, 10% dextran sulphate, 1% sodium dodecyl sulphate, 2×SSC, two DNA probes (2 ng/µl each) and 200 ng/µl salmon sperm DNA was predenaturated for 10 min at 75^°^ C and stabilised on ice for 10 min. Then, 40 µl of the hybridisation mixture was applied to each slide, and the slides were covered with plastic coverslips. The chromosomes and probes were denatured together in a thermocycler (PTC-100TM; MJ Research, Inc.) using a special adapter (The Slide Griddle^™^, Model SG96P; MJ Research, Inc.) for 7 min at 75^°^ C. The temperature was gradually decreased during hybridisation. Hybridisation was carried out overnight in a hybridisation oven (Biometra, OV3) at 37°C. The coverslips were removed and the slides were washed with 2×SSC for 2 min, 20% formamide in 0.1×SSC twice (5min each) and 2×SSC three times (8 min each). All the post-hybridisation washes were carried out at 42°C. The slides were incubated in 4×SSC/0.2% Tween 20 for 5 min at RT prior to detection, and then in 5% bovine serum albumin (BSA) in detection buffer for 15 min at 37°C. The probe labelled with digoxigenin was detected with fluorescein isothiocyanate (FITC)-conjugated sheep anti-digoxigenin antibody. Then, 50 µl of detection solution (final concentration of antibody 3 µg/ml) was applied to each slide and covered with a plastic coverslip. The slides were incubated in a moist chamber for 90 min at 37°C. Then, each slide was washed three times in 4×SSC/0.2 % Tween 20 at 42°C for 8 min, dehydrated through a series of ethanol solutions (70%, 85%, 96%; 1 min each) and finally air-dried. The slides were counterstained with DAPI (4’,6-diamidino-2-phenylindole, 2 µg/ml) or PI (propidium iodide) in antifade solution (AF1, Citifluor). According to Schwarzacher and Heslop-Harrison (2000), these conditions allow hybridisation between DNAs that share 85% sequence identity.

### Microscopy

The slides were examined under an Olympus BX-60 epifluorescence microscope (Hamburg, Germany), and images were taken with an Olympus E-520 camera (Olympus Imaging Europa GMBH, Hamburg, Germany).

## Results

### Cytogenetic disorders

Various cytogenetic disorders were observed in both amphiploid and its parental species. The hyperploid metaphases occurred at a frequency of 7.7% in the amphiploid and *A. longiglumis* compared to 0% in *Avena magna* H. C. Murphy & Terrell. The chromosomal bridges in anaphases (**Fig. 1b, f**) and telophases support the conclusion that the BFB cycle can occur in the free-nuclear syncytium. The frequency of bridges in *A. magna* and *A. longiglumis* was 52.38% and 25.0%, respectively. On the other hand, bridges were not observed in the amphiploid, while a low mitotic index was noted: only six out of 45 young caryopses showed anaphases and telophases. This indicates that analysis of more seeds could reveal the formation of bridges. The delayed single chromosomes or their groups were noted at a frequency of 7.7% in the amphiploid, and at 23.53% in *A. magna*. Chromosome elimination (Fig. 1b) was not observed in *A. longiglumis*. The delayed chromosomes or their fragments can form condensed micronuclei (Fig. 1a, c, e, g, h). In amphiploid and its parental species, micronuclei were noted in every syncytial endosperm. The maximum number of micronuclei per cell was 2 in the amphiploid and *A. magna*, while *A. longiglumis* showed single micronuclei. Micronuclei can also be formed by chromosomes or their fragments undergoing translocation (Fig. 1h). Nuclei with an irregular shape and showing different level of condensation were observed in 5.88% of *A. magna*, and 15.38% of *A. longiglumis* caryopses (Fig. 1d, e).

**Fig 1.**
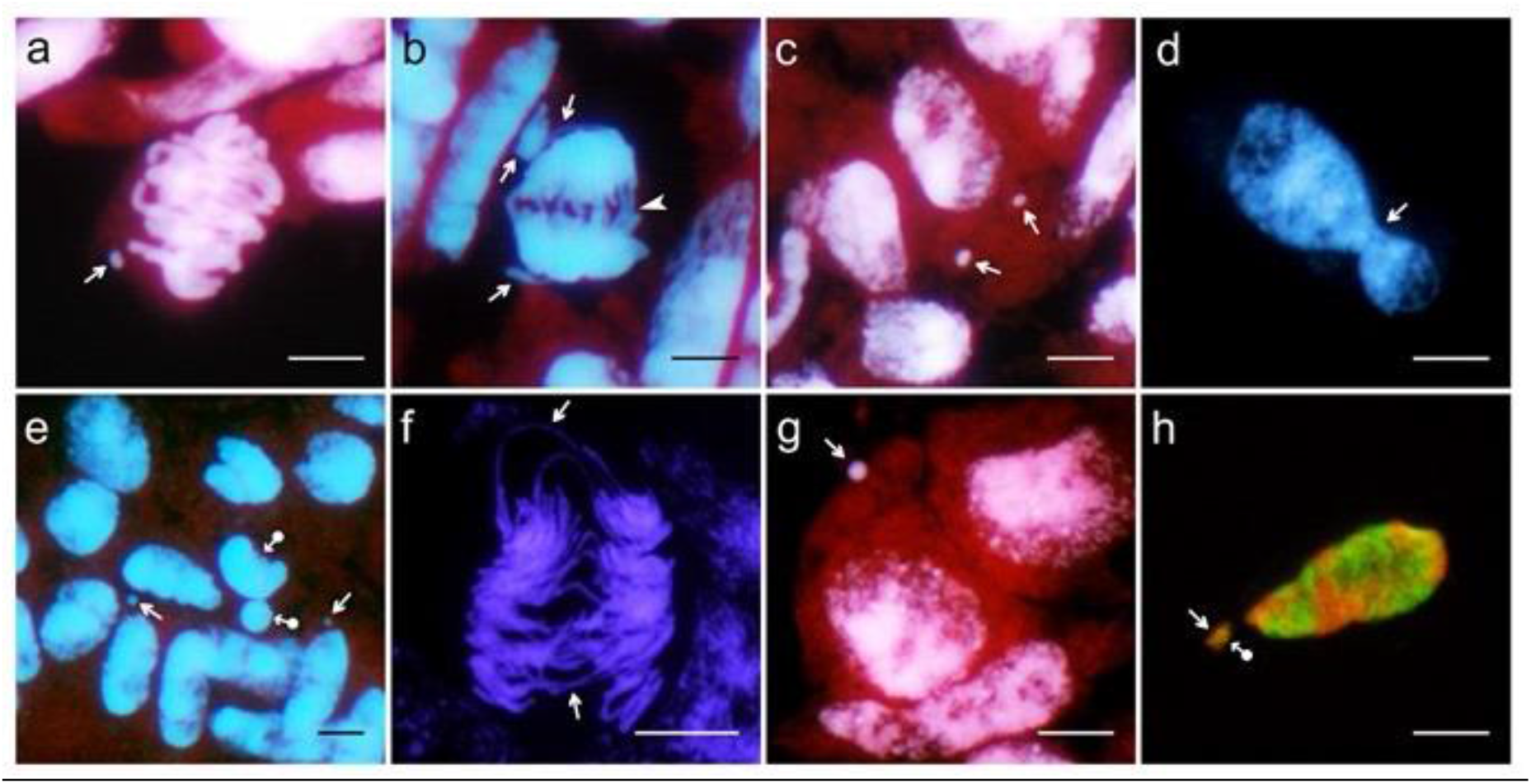
Examples of cytogenetic disorders in: *A. maroccana* (**a-c**), *A. longiglumis* (**d-f**) and *A. maroccana* × *A. longiglumis* amphiploid (**g** and **h**). **a** – metaphase with a fragment of chromosome forming a micronuclei (arrow), **b** – anaphase with multiple bridges (arrowhead) and two precocious chromosome units at the poles (arrows) which are still connected to the poles, **c** – nucleus with two micronuclei (arrows), **d** – an irregular nucleus (arrow) before the creation of two nuclei of different sizes, **e** – nuclei with micronuclei (arrows) and a highly condensed nucleus extruding a micronucleus of the same kind (dot-arrows), **f** – anaphase with several bridges (arrows), **g** – a micronucleus (arrow) and a group of nuclei at the early prophase, **h** – a micronucleus composed of two genomes, red C genome (arrow) and green A genome (dot-arrow). DAPI fluorescence for **a** – **g** and counterstained by propidium iodide for **a, b, c, e, g**. For **h** - red fluorescence for *A. eriantha* probe (C genome) and green fluorescence for *A. nuda* probe (A genome). Scale bars 10 µm

The expected number of chromosomes in endosperm (**Table 1**) and genomic formulae are as follows: amphiploid 63, AAACCCAlAlAl; *A. maroccana* 42, AAACCC; and *A. longiglumis* 21, AlAlAl. The observed disorders, especially the elimination of chromosomes and their fragments as well as the micronuclei formation, gave rise to new sets of chromosomes and the series of chromosome numbers (Table 1). A narrow variation was noted in this characteristic in *A. maroccana*, whereas in the amphiploid and in *A. longiglumis* the series of the chromosome numbers exhibited a broad variation. The expected number of chromosomes typical for the endosperm tissue (3*n*) was observed in the amphiploid and both parents. The modal number of chromosomes was distinctly lower and oscillated at 2*n*. The DNA amplification typical of the hyperploidy level occurred in the amphiploid as well as the paternal species.

**Table 1.**
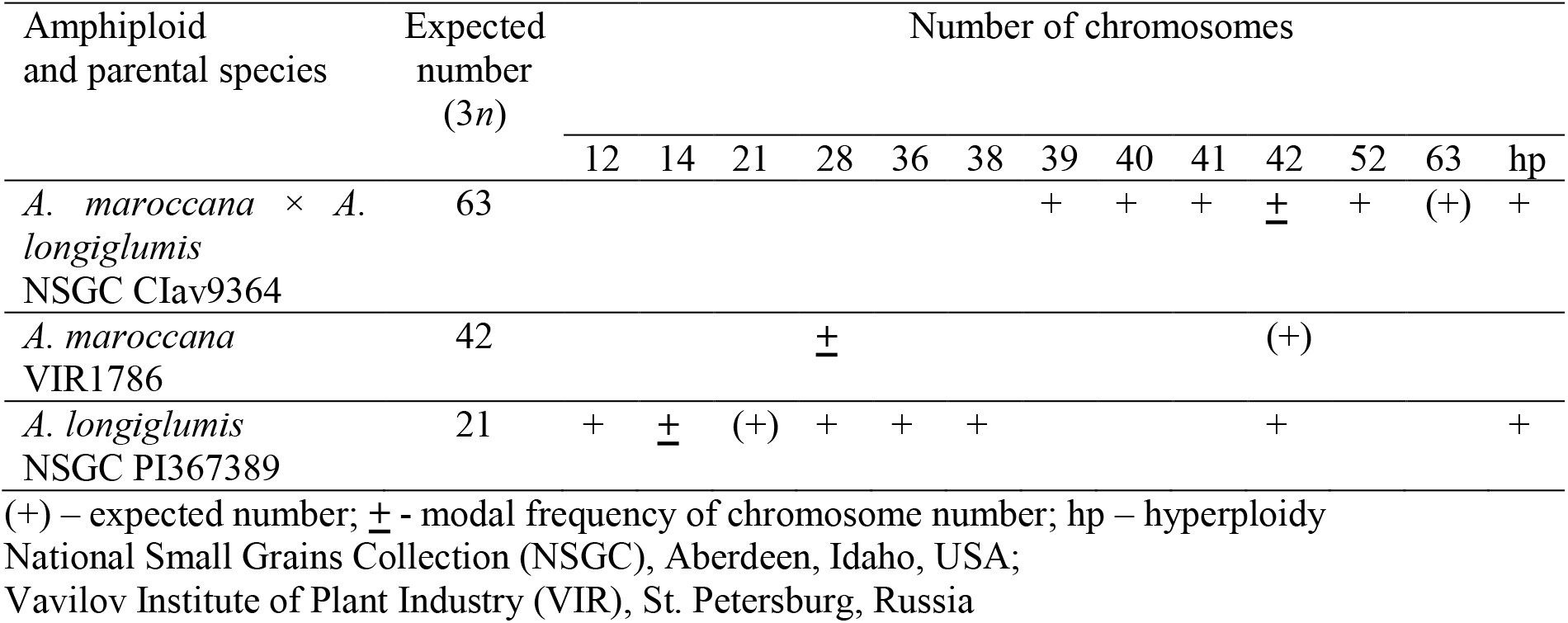
Variation of the chromosome number in the free-nuclear endosperm of the amphiploid and its parental species

### GISH variation

#### Translocations in A. marrocana

The two genomic probes obtained from *A. nuda* (AsAs genomes, green fluorescence) and *A. eriantha* (CpCp genomes, red fluorescence) were used for the detection of A and C genomes in the endosperm nuclei of *A. maroccana* (**Fig. 2**). Six C/A and four A/C translocations were noted in the prophase nucleus (*2n = 28*, Fig. 2a). At least one pair (1, 1’) were tiny and showed equal reciprocal terminal translocations. The two C/A translocations in the upper right part overlapped partially. The lower one, marked with an arrowhead, ended with the terminal green points of the A genome. In anaphase, the translocations in the daughter chromosome groups appeared the same (2*n* = 28, Fig. 2c), and four C/A and two A/C translocations were observed. Terminal translocations dominated in the prophase nucleus, of which two were intercalary. Even in the anaphase groups, the terminal ones dominate (Fig. 2c). The translocations shown in Fig. 2c can be better differentiated by observing **Fig. S1** (on-line supplement, Additional File S1). In addition to reciprocal translocations (Fig. 2a), single and non-reciprocal translocations were also noted. The differences in the number of both types of translocations proved that the translocation status of the endosperm nuclei in the ‘maroccana’ species is variable. Here, one should consider the possible elimination of translocated segments in the micronuclei (Fig. 1h for the amphiploid). The DAPI image of the prophase overlapped by the green fluorescence of the AA genomes shown in Fig. 2b indicates that the CC genomes are not homologous to the genomic DNA from *A. nuda* and that A genome of *A. maroccana* is homologous to the As genome from the *A. strigosa* group (Fig. 2b).

**Fig 2.**
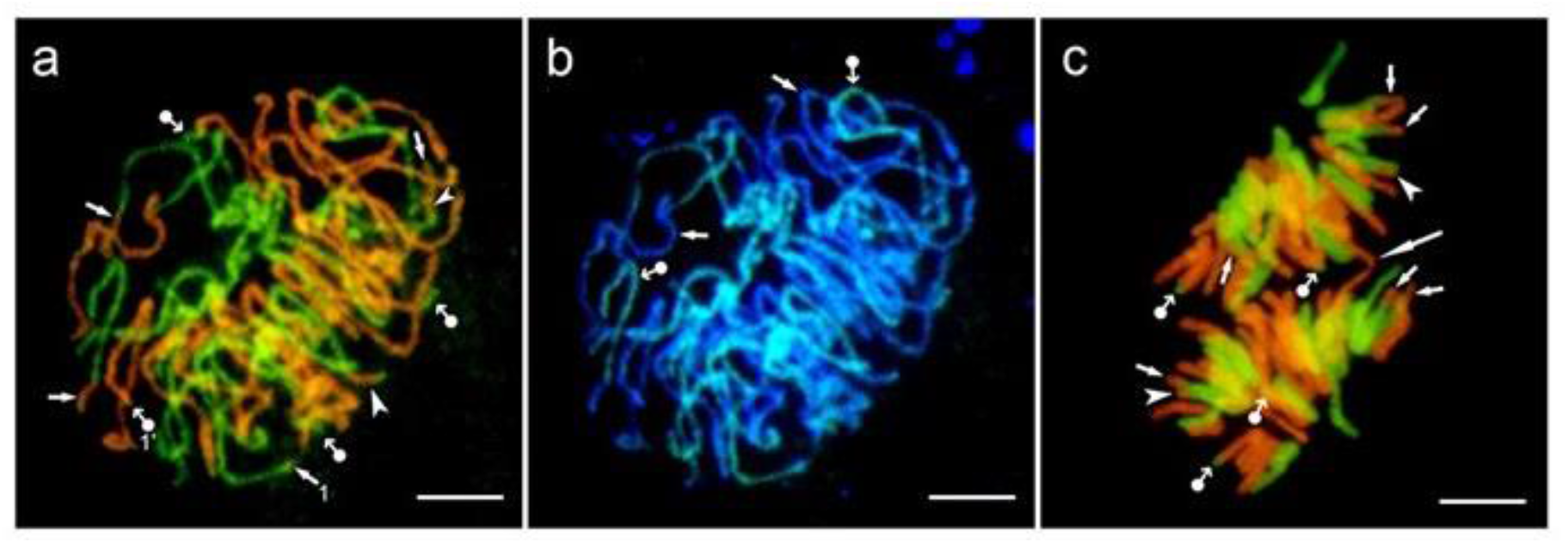
Chromosome rearrangements in *A. maroccana* detected by GISH. **a** – prophase with four A/C (dot-arrows) and six C/A translocations (arrows for terminal, arrowheads for intercalary), 1 and 1’ – a pair of terminal reciprocal microtranslocations; **b** – green chromosomes of the A genome marked with the *A. nuda* probe (dot-arrows) and DAPI blue chromosomes of the C genome (arrows), **c** – anaphase translocations (small arrows) and a bridge (large arrow); in both groups of chromosomes, the same number of translocations is marked by arrows for terminal C/A, arrowheads for subterminal C/A and dot-arrows for A/C. Scale bars 10 µm

For a sample of nuclei analysed (*n* = 15), the arithmetic average of the C/A translocations was 5.47 per nucleus, while that of the A/C translocations was 2.53. The first type of translocation was less variable (coefficient of variation V = 16.8%, min-max values 4-6, modal number 6) than the second (coefficient of variation V = 36.1%, min-max values 2-4, modal number 2). For both types of translocations, the distribution curves were leptokurtic. A bimodal distribution was observed when both curves were overlapped (min A/C and max C/A). Terminal translocations dominated in both C/A and A/C types; however, in the first type subterminal translocations and in the second microtranslocations were also noted.

#### Translocations in the amphiploid

The DAPI fluorescence and two genomic probes obtained from *A. nuda* (AsAs genomes, green fluorescence) and *A. eriantha* (CpCp genomes, red fluorescence) allowed detecting the A and C genomes and chromosome rearrangements in the endosperm nuclei of the amphiploid. The DAPI staining enabled clear discrimination of individual chromosomes and the level of their chromatin condensation (**Fig. 3a**). Chromatin condensation was even along the chromosomes a-e and chromosome 3, but not so in chromosomes 1 and 2, in which the terminal sectors or arms were less condensed. The use of the Cp probe for genome C (Fig. 3b) revealed that chromosomes a-e belonged to the C genome and b, c and d chromosomes showed weaker hybridisation signals on the large terminal sectors. In the presented metaphase, eight terminal and one intercalary C/A translocation were found. There were no reciprocal translocations.

**Fig 3.**
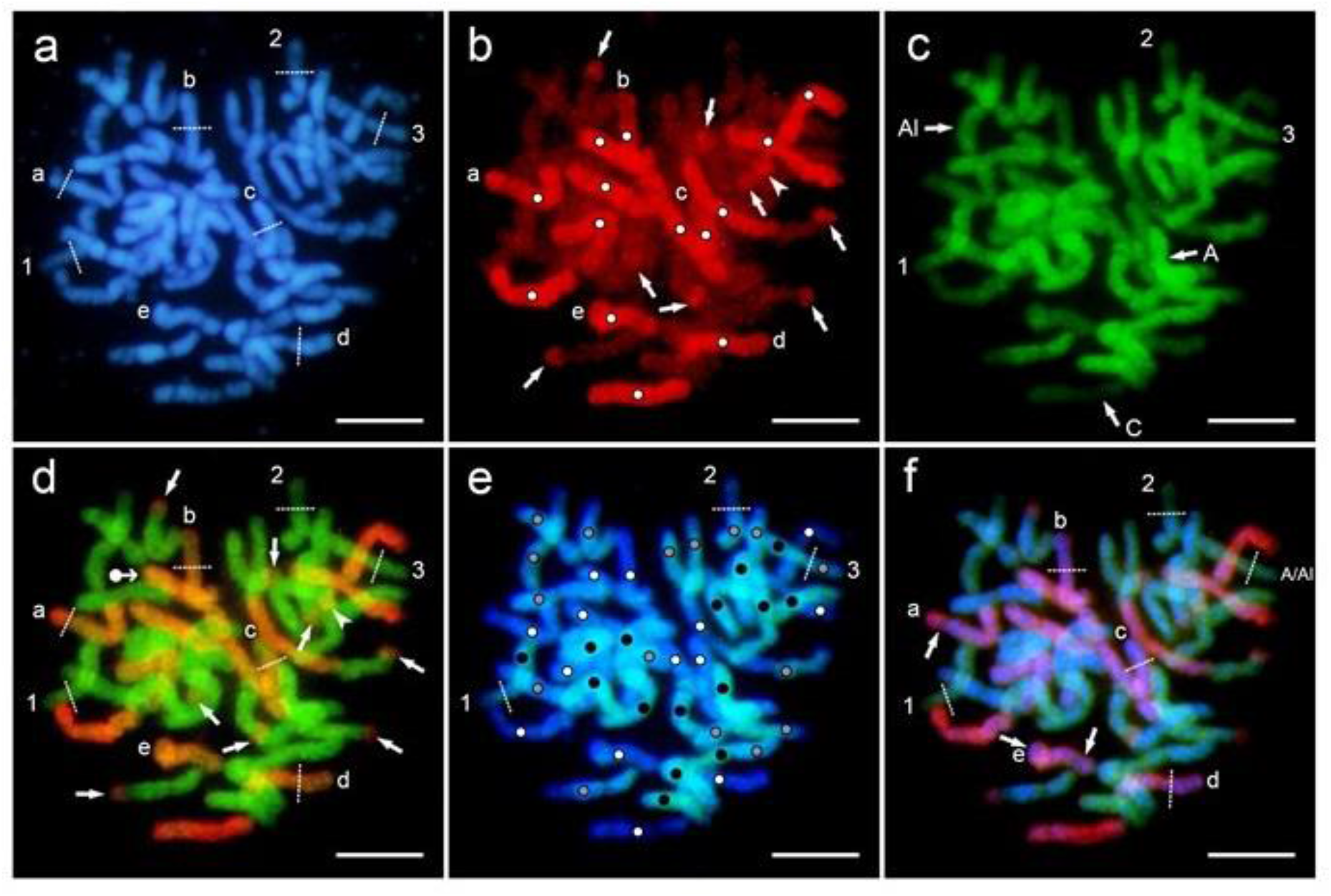
Detection of genomes and translocations in the Am×Al free-nuclear endosperm. Two probes of the total genomic DNA were used - Cp genome from *A. eriantha* (red) and As genome from *A. nuda* (green); in addition, the slides were counterstained by DAPI (blue). **a** – a metaphase (3*n* = 42) showing DAPI fluorescence, with different sectors in chromosomes separated by dotted lines, numbers and letters; (1 and 2 mark the sectors of lower DNA condensation, and for others, designation of sectors with reduced homology to the probes used (see **b**-**f**); **b** – red fluorescence of eight terminal (arrows) and one intercalary (arrowhead) C/A,Al translocations marked by using the *A. eriantha* probe, with white dots indicating the C genome chromosomes; **c** – green fluorescence shows the homology of three genomes (C, A, Al) of the amphiploid after hybridisation of the *A. nuda* probe; **d** - red and green fluorescence of two overlapped probes marks the same translocations as in **b**, and in addition one tiny terminal A/C translocation (dot-arrow); for chromosomes ‘a’ to ‘e’, the sectors of lowered homology to the C genome probe or even arm translocations (see ‘c’ chromosome) are shown; **e** – the metaphase marked by DAPI and green fluorescence of the *A. eriantha* probe – three genomes can be identified: white dots for C, black for A and grey for Al genome; for chromosomes ‘1’ and ‘2’ arms of lower DNA condensation and for chromosome ‘3’ the possible A/Al arm translocation (also shown in Fig. 3f) are distinguished; **f** – the metaphase with the DAPI fluorescence and fluorescence of both probes overlapped (for chromosomes ‘a’ and, ‘e’ intercalary and for chromosomes ‘b, ‘c’ and ‘d’ large terminal lowering of homology to the Cp probe or chromosome rearrangements are well visible). Scale bars 10 µm

Labelling with the As probe of the A genome resulted in the distinct fluorescence of the A genome chromosomes in the amphiploid (Fig. 3c). The C genome chromosomes showed a weak fluorescence. In addition, the chromosomes with an intermediate fluorescence belonging to the Al genome were identified. This would prove the higher chromosome homeology between the A and Al genomes. The difference between these two genomes in the probe binding was quantitative, and thus, it was difficult to discriminate them. Chromosomes 1-3 belonged to the A and A/Al genomes, of which chromosomes 1 and 2 showed a weaker fluorescence, due to lower chromatin condensation (Fig. 3a). Chromosome 3 had a weaker hybridisation signal on its right arm having the homogenous condensation of chromatin (Fig. 3a). The C/A translocations in Fig. 3b were better discriminated after the fluorescence of both probes labelling the C and A genomes was overlapped. In addition, a small A/C terminal translocation was detected (Fig. 3d). The dotted lines on chromosomes a-e of the C genome having the homogenous chromatin condensation (Fig. 3a) indicate large fragments along the chromosome with various level of homology to the probe used. The superposition of the fluorescence of *A. nuda* probe (As genome) on the DAPI fluorescence enabled the discrimination of chromosomes belonging to the A, C and Al genomes in the amphiploid endosperm (Fig. 3e). The dotted lines also indicate a weaker binding of the probe in chromosomes 1 and 2 which correlated with a lower level of chromatin condensation (Fig. 3a) and possible translocation of the entire arm between the A and Al genomes in chromosome 3. Differentiation in the level of homology in the C chromosomes with respect to the Cp genome probe is shown in Fig. 3f with combined DAPI fluorescence and fluorescence of both probes (chromosomes a, b, c, d, e). Large, continuous fragments of chromosomes with a weaker probe binding may also demonstrate intragenomic rearrangements.

For a sample of nuclei analysed, the arithmetic average of the C/A translocations was 4.14 per nucleus, while that of the A/C translocations was 1.86. The first type of translocation was less variable (coefficient of variation V = 57%, min-max values 0-8) than the second type (coefficient of variation V = 70%, min-max values 0-4). The total number of translocations ranged from 0 to 11. For a small sample (*n* = 13), the distribution curve was platykurtic for the C/A translocations with modal numbers 2, 3, 4, 6 and 7, while it was leptokurtic for the A/C translocations with modal number 2. The variation of translocations was distinctly larger in the amphiploid compared to the maternal species.

#### Spatial arrangement of genomes

At the interphase and various stages of mitosis, the chromatin and chromosomes were not distributed randomly. A prophase nucleus in *A. maroccana* showed a sectorial-concentric domain arrangement of the A and C chromosomes (**Fig. 4a**). Terminal and intercalary translocations were located at the periphery. In the amphiploid, the central part of a nucleus (marked by a dot-arrow in Fig. 4b) was found to be more condensed than its periphery (marked by arrows) and related to the green A genome. The outer part showing a weak green fluorescence indicated the location of the Al genome; however, another interpretation is also possible; the outerpart might indicate decondensed chromatin of the A and Al genomes. The red C genomes were organised in the form of bands while in the outer part of the nucleus they were condensed (marked by white dots). In addition, a small dot of the C genome (small arrow) was seen in an area of the Al or A + Al genomes. This fluorescence signal can be interpreted as a chromosome microrearrangement. Probably, such a small change on metaphase chromosomes cannot be detected. The green A and Al genomes (indicated by letter a in Fig. 4c) *versus* the red C genomes (letter b) appeared to be situated side-by-side. The composite fluorescence red, green and blue (Fig. 4d) revealed a new arrangement of genomes. The red Cp and green A genomes dominated the outside area of the prophase nucleus(dots and arrows in Fig. 4d), while the blue Al genomes also having a weak green fluorescence were situated mainly inside and only tiny parts of Al chromosomes were located outside of the prophase nucleus (asterisks in Fig. 4d). The arrangement of chromosomes in Fig. 4d can be described as a sectorial-concentric pattern, which is similar to that observed in *A. maroccana*.

**Fig 4.**
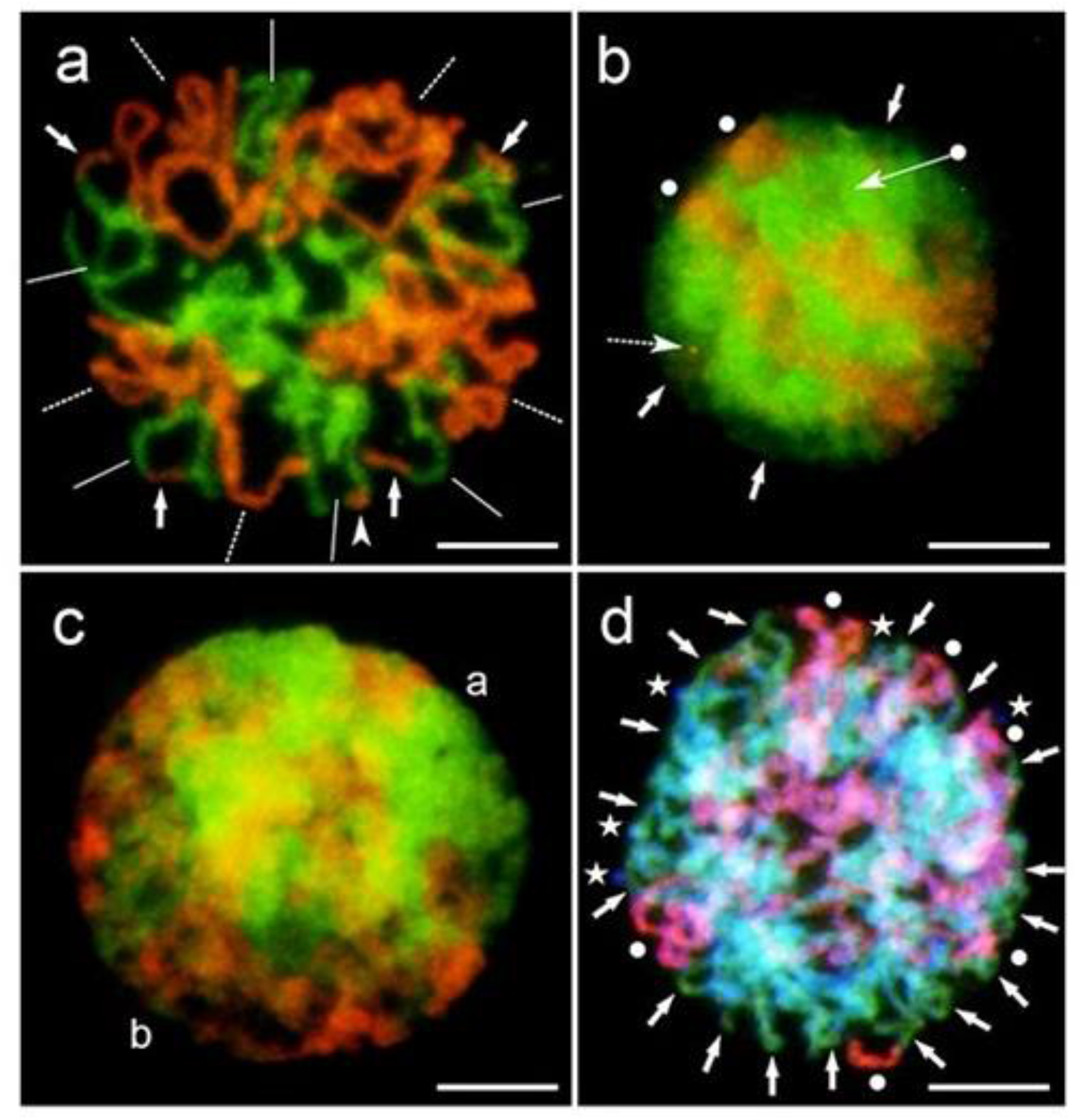
Spatial arrangement of genomes in *A. maroccana* (**a**) and the amphiploid (**b**-**d**). **a** – a prophase sectorial-concentric distribution of the A (lines) and C (dashed lines) genome chromosomes, with terminal (arrows) and intercalary (arrowhead) translocations found outside; **b** – an interphase-prophase nucleus with a condensed centre (dot-arrow), decondensed external parts (arrows), the C genome condensed at the periphery of the nucleus (white dots) and an orange dot showing the A/C chromosome microrearrangement (small arrow with dashed line); **c** – a side-by-side arrangement of the green A,Al genome (a) and red C genome (b) in the nucleus of an early prophase; **d** – a sectorial-concentric arrangement of genomes in the nucleus of prophase (green A genome (arrows), red C genome (white dots) and blue Al genome (asterisks)). Scale bars 10 µm

## Discussion

### Cytogenetic disorders

Combining the genomes of different species, for instance wheat and rye, leads to occurrence of chromatid or chromosomal bridges in the hybrid mitotic cycle, and the BFB cycle (Lukaszewski 1995). The BFB cycle was also noted in the cells of maize callus culture. It appeared at heterochromatic knobs, and the broken ends were later healed by telomere sequences (Santos-Serejo and Aguiar-Perecin 2016). In barley and maize, Kilian et al. (1998) noted a decrease in telomerase activity during endosperm development as well as telomere shortening. In human cancer cells, telomere loss activates the BFB cycle and consequently leads to many chromosomal rearrangements, including micro-translocations and deletions (Lo et al. 2002). Such events are supported by data on telomerase inactivity in *Arabidopsis*, which was shown to increase fusion and breakage of chromosomes at anaphase-telophase and subsequently leads to unequal distribution of chromatin and rDNA to sister nuclei (Siroky et al 2003). This cytogenetic behaviour was multiplied during repeated BFB cycles.

In a syncytial endosperm of Am×Al, no bridges were noted, but were observed in its root tissues (Świetlikowska 2008). In the Triticale endosperm, Kaltsikes et al. (1975) observed a low proportion of bridges and attributed such a frequency to the shorter anaphase-telophase stage. In the other amphiploids of oats, bridges were formed together with rings and telocentrics (Tomaszewska and Kosina 2018). These events were evidently components of the BFB cycle occurring in the free-nuclear stadium of the changing oat embryo sac. In a group of oat amphiploids and their parental species, the frequency of bridges was determined at a similar level, but the sum of various cytogenetic anomalies was found to be comparatively much higher in amphiploids. (Tomaszewska and Kosina 2018).

Elimination of chromosomes and their fragments and micronuclei formation are two events which are important for the subsequent development of endosperm tissue. In *A. sativa* varieties, elimination of micronuclei from microspores in the form of microcytes was observed, which lowered the male fertility (Baptista-Giacomelli et al. 2000). Other data showed that pearl millet-origin micronuclei of the wheat × pearl millet hybrid embryos were extruded from the interphase nuclei in the form of highly heterochromatinised bodies, and were finally fragmented (Gernand et al. (2005). Highly condensed irregular nuclei of various sizes showed characteristics of apoptosis. Such type of nuclear degeneration during programmed cell death has been noted for antipodals in wheat (An and You 2004). It is highly probable that the high frequency of micronuclei in Am×Al and its parents is responsible for the selective elimination of anomalous structures with DNA and reduction in the level of syncytial ploidy.

A consequence of cytogenetic anomalies in the endosperm is the formation of dysfunctional nuclei and their elimination during apoptosis. Kaltsikes et al. (1975) suggested that in Triticale, irregular endosperm nuclei having different sizes and amounts of DNA were formed after breaks of bridges. In addition, many nuclei were noted to be necrotic (apoptotic?), and undoubtedly were selectively eliminated. Such a selection has also been reported in an oat amphiploid, *Avena barbata* Pott ex Link × *A. sativa* (Kosina and Tomaszewska 2011), as well as in some oat species (Tomaszewska and Kosina 2018). The apoptotic selection of nuclei can decrease the level of endosperm ploidy and maintain a lower modal number of chromosomes than that expected for a triploid tissue. This phenomenon not only occurs in polyploid forms but has also been observed in a diploid species (e.g. *A. longiglumis*, Table 1) and is probably necessary to stabilise development of the storage tissue.

### Translocations

The observed translocations occurred between all oat genomes in the amphiploid nuclear endosperm. These chromosomal events were predominantly terminal and were of various sizes. In the endosperm, translocations can be accompanied by other structural rearrangements, including the BFB cycle. In addition, somatic (mitotic) crossing over (SCO) may rarely occur in the endosperm tissue at each developmental stage (Tomaszewska and Kosina 2018). SCO could be observed as a mosaic of GISH signals (+ or -) across the chromosome, and this uncommon event cannot be neglected in the endosperm.

A dotted pattern of probe fluorescence, especially red *versus* blue, is shown in Fig. 3f. Such a pattern is interpreted as weak hybridisation of the genomic DNA probe at a low-homologous target (Schwarzacher and Heslop-Harrison 2000). This sequence-homology-dependent strength of fluorescence has been well exemplified by Seijo et al. (2007) in their study, in which the genomic DNAs obtained from several wild species of peanut were applied as probes to detect the chromosomes of *Arachis hypogaea* L.. The fluorescence of dotted scattered signals was observed to differ with their intensity and number depending on the homology of the probe to the genome of *A. hypogaea*. In the amphiploid, weak hybridisation signals can be located at the parts of chromosome showing less condensed chromatin. Such parts were visible as large terminals (Fig. 3a – DAPI fluorescence, chromosomes 1 and 2). Thus, in the oat amphiploid, the segments of chromosomes with no visible fluorescence signals of a given genomic DNA probe can be considered as fragments having a reduced homology to the applied probe. These segments do not have sharp limits due to the dotted hybridisation at the contact with parts having uniform fluorescence.

The heterochromatic regions of chromosomes have been recognised as more fragile than the euchromatic sites (Jannsen et al. 2018). In the case of *A. sativa*, most chromosomal exchanges occur in the heterochromatic C genome (Chen and Armstrong 1994; Jellen et al. 1994). Such a tendency to create breaks and chromosome fragments was also exemplified for the C genome of *Avena fatua* L. and other oat hexaploids by Yang et al. (1999). Fluminhan and Kameya (1997) discovered that breaks were associated with heterochromatic knobs in maize chromosomes. C-banding and *in situ* DNA hybridisation are two useful techniques for studying the interrelations between translocations and heterochromatin. The compatibility of heterochromatic C-bands with the heterochromatin repetitive sequences at the *in situ* hybridisation site has been demonstrated for some plants, including *Allium fistulosum* L. (Irifune et al. 1995) and *Alstroemeria* L. (Kuipers et al. 1997). Thus, the frequency of translocations in the oat endosperm must be related to the differences in the amount of heterochromatin between the genomes. In the amphiploid and its maternal species, the dominance of the C/A-type translocation is distinct at terminal positions. The dominance of terminal translocations in the oat chromosomes is corroborated by the diversity of repeat sequences localised in them (Liu et al. 2019). Compared to other oat tissues, the dose of heterochromatin is increased in endosperm due to its input in the CCC maternal genomes. Such a relation was also proved for the endosperm of *Arabidopsis* (Baroux et al. 2007). Thus, a positive correlation between the dose of heterochromatin and the C/A translocation number in the endosperm of the amphiploid can be expected. The comparative numbers of translocations in root mitoses have not been studied for the amphiploid.

Using a probe of *A. eriantha* (Cp genome), Nikoloudakis and Katsiotis (2015) determined four major C/A terminal translocations in the root tissue of a triploid hybrid *A. magna* × *A. longiglumis*. In *A. maroccana* (synonym *A. magna*), the number of such translocations was determined as six by using oligo-probes (Fominaya et al. 1995; Luo et al. 2018b) and as eight by using genomic DNA (Leggett et al. 1994) or oligo-Am1 probe (Luo et al. 2018a). The translocations were tiny and terminal or sub-terminal. Leggett et al. (1994) concluded in their study that non-reciprocal C/A translocations dominated. These data prove that the translocations occurred exclusively between the C and A genomes belonging to *A. maroccana* (*A. magna*), while the chromosomes of *A. longiglumis* did not participate in these exchanges in the triploid hybrid. However, the amphiploid endosperm differs in this respect. Fig. 3e confirms the occurrence of C/A and C/Al translocations. The first ones are older and also noted in *A. maroccana*, whereas the second ones are new and occurred during the hybridisation process. Furthermore, the A/Al translocation should be treated as a rare and new event (Fig. 3f).

In an *A. strigosa* × *A. maroccana* amphiploid (2*n* = 42), eight C/A translocations (six terminal and two intercalary) and two tiny terminal A/C translocations were observed in the root tissue (Ueno and Morikawa 2007). These results are very similar to those reported in this study on young nuclear endosperm (Fig. 3); however, the number of translocations can increase with the chromosome number (Table 1). According to Hayasaki et al. (2000), the number of translocations in oats is positively correlated with the level of ploidy.

The above amphiploids differ from each other in the following: the introduced genomes (As or Al); the parental status of species (maternal or paternal), and the cytogenetic stability of the tested tissues (root or endosperm). Despite these differences, the high similarity of the results proves the dominant role of the C genome in the translocations occurring of in both tissue types -root and endosperm. The data show that the C/A and A/C translocations vary by number in the same nucleus, and are therefore non-reciprocal. A question arises whether this is a natural state or whether the reciprocal translocations are eliminated. Schubert and Lysak (2011) postulated that non-reciprocal translocations are, in fact, a result of unbalanced chromosome segregation. In the syncytial endosperm, the selection of dysfunctional nuclei and micronuclei (Fig. 1e), as a product of such segregation, occurs, and the translocated chromosomes or their fragments (Fig. 1h) can be removed, following which the level of ploidy decreases (Table 1). However, the unequal number of C/A *versus* A/C translocations in the *A. maroccana* anaphase is maintained in the sister nuclei (Fig. 2c), which does not support the conclusion about the unbalanced segregation origin of non-reciprocal translocations. The same ratio of both translocations observed in root tissue (Leggett et al. 1994; Hayasaki et al. 2000; Nikoloudakis and Katsiotis 2005; Ueno and Morikawa 2007) proves that such translocations are oat-specific and truly non-reciprocal. The present study shows that reciprocal translocations are not frequent.

Recent studies have indicated a special role for *A. longiglumis* in the evolution of oat tetra- and hexaploids. The Al genome present in the amphiploid is considered ancestral to the other A genomes (Holden 1966; Drossou et al. 2004). Fu (2018) attributes a key role to *A. longiglumis* in the evolution of oats with the AACC and AACCDD genomes. Analyses of cp and mt genomes proved that *A. longiglumis* was a maternal component that created *A. insularis* (AACC) and subsequently a paternal species with *A. insularis* as a maternal species in the evolution of the AACCDD species.

When crossed with *A. maroccana, A. longiglumis* (AlAl genomes) showed a higher level of meiotic chromosome homology compared to *A. strigosa* (AsAs genomes) (Rajhathy 1991). In this species, the accession Cw 57, which was used to create the amphiploid (Am×Al is a young breeding form, created in the early 1980s; Ref.: https://), increases the homeologous pairing in hybrids (Rajhathy and Thomas 1972; Thomas and Al-Ansari 1980). According to Jellen and Leggett (2006), the A-genome of *A. maroccana* differs from that of the diploid species, and so the genomic formula of this species should be rather CCDD, and not AACC. This was also supported by Yan et al. (2016). The present paper shows that the hybridisation signal obtained for the *A. nuda* probe (AsAs group of diploids) is clear and uniform along AA chromosomes in both *A. maroccana* and Am×Al, while it is weaker in Am×Al for the ‘longiglumis’-chromosomes. Such data seem to be contradictory to those cited by Jellen and Leggett (2006). However, assuming that the A and D genomes are evolutionarily younger than the Al genome, the strong hybridisation signal between the *A. nuda* probe and the amphiploid AA genomes, and weaker signal observed for the Al genome, can prove the similarity between A and D genomes as shown by Linares et al. (1998). The fluorescence *in situ* hybridization analyses of the root mitoses which were recently carried out in *A. maroccana* and *Avena murphyi* Ladiz. using specific probes for the C and D genomes confirm the genome composition CCDD in tetraploids previously designated as AACC (P. Tomaszewska, pers. comm.). Considering the role played by *A. longiglumis* in the evolution of AACC and AACCDD oats and its phylogenetic closeness to them (Chew et al. 2016; Yan et al. 2016; Fu 2018), the genomic formula of the amphiploid can be accepted as CCDDAlAl and that of its endosperm as CCCDDDAlAlAl.

The old, large translocations were introduced by *A. maroccana*, while the new ones may have arisen between the ‘maroccana’ AACC (CCDD) genomes and the new Al genome. The A/Al translocation should considered as new (Fig. 3f). As a rule, meiosis in parental species having old translocations is regular, but with defects in hybrids and amphiploids (Holden 1966; Rajhathy 1971; Ladizinsky 1998; Leggett 1998), which can be attributed to the occurrence of new translocations and other chromosomal rearrangements. These cytogenetic events can be more frequently observed in the endosperm than in root meristems; however, the selection of dysfunctional nuclei decreases the number of chromosomes and their rearrangements that could be observed in the endosperm (Table 1; Tomaszewska 2017). This type of selection is commonly made through the apoptosis process (Tomaszewska and Kosina 2018).

### Spatial arrangement of genomes

The non-random distribution of chromosomes in the metaphases was documented in an earlier study by Müller (1909) in the root cells of *Yucca* L.. This observation was later supplemented by the findings reported for other Monocotyledonae species by Darlington (1956). Two types of chromosomes, tiny and large, which were separately scattered, were identified. This phenomenon was clearly seen in hybrids in which two different sets of chromosomes of parental origin formed a nucleus. In common, chromosomes are arranged either side-by-side or concentrically (Bennett 1984). Chromosome separation can be identified by the difference in dimensions, as well as in C-banding, as reported for the hybrid *Hordeum* L. × *Psathyrostachys* Nevski ex Roshev. (Linde-Laursen and Jensen 1991). In barley, shorter and more heterochromatic C-banding chromosomes were found in the centre during metaphase, while in the case of *Psathyrostachys*, longer chromosomes were observed concentrically outside. The concentric pattern can often be realised as a sectorial arrangement in which the size of the sectors corresponds exactly to the number of chromosomes or the amount of DNA in the separated genomes (Kosina and Heslop-Harrison 1996; Kosina 1999). Leitch et al. (1991) proved that the spatial separation of chromatin, and then chromosomes, from the genomes of different origins persists throughout the cell cycle.

In this study, the sectorial-concentric arrangement of chromatin is exemplified for *A. marroccana* and Am×Al (Fig. 4). At interphase, a side-by-side pattern was also noted. The outer parts of the interphase nucleus showed a weaker green fluorescence of chromatin which can be interpreted as Al genome. In this position, it is possible to locate nucleoli non-hybridising with the DNA of a probe and showing a weak fluorescence. However, Heslop-Harrison et al (1993) proved that the nucleoli in somatic interphase nuclei were located in the centre of the nucleus, and so the weak fluorescence of peripheral chromatin can indicate some decondensation of the Al chromatin. This conclusion is confirmed by the location of 25S rDNA loci which was not outside but in the centre of the nuclei in the oat amphiploid (Tomaszewska 2017). The pattern of genome arrangement is sectorial-concentric in the prophase nuclei, but in interphase it is different. Gleba et al. (1987) proved that in old somatic distant hybrids the genome arrangement changed from segmental to sectorial. In the oat endosperm, nuclei approaching apoptosis showed concentric arrangement of genomes (Tomaszewska and Kosina 2013). It seems probable that in developmentally unstable hybrid tissues, such as endosperm or cell cultures, the pattern of genome arrangement can be variable.

## Concluding remarks

Among the cytogenetic disorders, appearance of bridges and elimination of chromatin in the final form of micronuclei, including translocated segments, are considered important for the functioning of the future endosperm tissue. A series of disorders can be associated with the BFB cycle. A decrease of free-nuclear endosperm ploidy from the level of 3*n* and higher proves that many polyploid nuclei are eliminated during apoptosis. The modal number of chromosomes is close to 2*n* and seems to be optimal for endosperm development in both parental forms and amphiploid. Structural specificity of the subtelomeric regions of *A. maroccana* chromosomes which are in the form of heterochromatin blocks facilitates the translocation of its C genome fragments into other oat genomes. The maternal species of the Am×Al amphiploid has its own large C/A translocations, and in the free-nuclear endosperm, these translocations occur similarly as in root tissues, compared with the reference data. Such events relate to the nuclei in both tissues with the same or similar level of ploidy.

C/A translocations occurred more commonly than A/C translocations and usually in the terminal parts of chromosomes, while subterminal or intercalary translocations occurred less often. Such status seems to be conservative. Generally, translocations were single and non-reciprocal, or rarely reciprocal. The occurrence of A/C translocations was more variable than the C/A type. Small or even microrearrangements of chromosomes were also observed in the amphiploid. Large fragments in the C chromosomes that showed poor homology to the used probe may prove that the C genome underwent intragenomic rearrangements, which seems possible with its heterochromatin status. During the microevolution of the amphiploid, new rearrangements took place invoving the Al genome, which were noted as a broader variation of the number of translocations than in *A. maroccana*. The last reference data prove that *A. longiglumis* is a possible progenitor of AACC and AACCDD oats. Both A (As) and Al genomes studied in the present paper showed a high homeology, so it is proposed that the genomic designations of AACC tetraploids should be changed into CCDD or CCDlDl (Dl – the *A. longiglumis* genome changed during the evolution of the AACC tertraploids). Such an approach needs more research focused on the level of homeology/homology between the A genome in AACC and the variants of the Al genome.

It has also been proved that the parental genomes of the Am×Al amphiploid are not mixed together but remain spatially separated and arranged in a sectorial-concentric pattern. Furthermore, a side-by-side pattern was observed for this taxon, especially in the early prophase. Thus, the pattern of the spatial genome arrangement appears to be variable, particularly in less stable tissues such as the endosperm of hybrids or plant tissue cultures.

## Compliance with ethical standards

### Ethics approval

The authors guarantee compliance with ethical standards.

### Consent to participate

Not applicable

### Consent for publication

Not applicable

### Availability of data and material

Raw cytogenetic pictures stored electronically.

### Code availability

Not applicable

#### Abbreviations

A file included

## Authors’ contributions

PT designed and conducted all cytogenetic analyses and interpreted the results. PT and RK prepared figures and wrote the article. RK provided research idea, supervised the experiments and research documentation. The authors read and approved the manuscript.

## Acknowledgements

The authors would like to thank the following institutions for providing seed samples of oat amphiploid and species: National Small Grains Collection (Aberdeen, Idaho, USA) and Vavilov Institute of Plant Industry (St. Petersburg, Russia). This research was financially supported by the statutory fund of the Institute of Experimental Biology, University of Wroclaw, Poland. This report is a part of the PhD thesis written by P. Tomaszewska at the University of Wroclaw, Poland.

## Electronic supplementary material

The online version of this article (https://......) contains supplementary material Fig. S1 related to Fig. 2c in the article, which facilitates the detection of the translocations in the analysed anaphase. The supplementary figure is available to authorized users.

## Supplementary file

**Fig S1.**
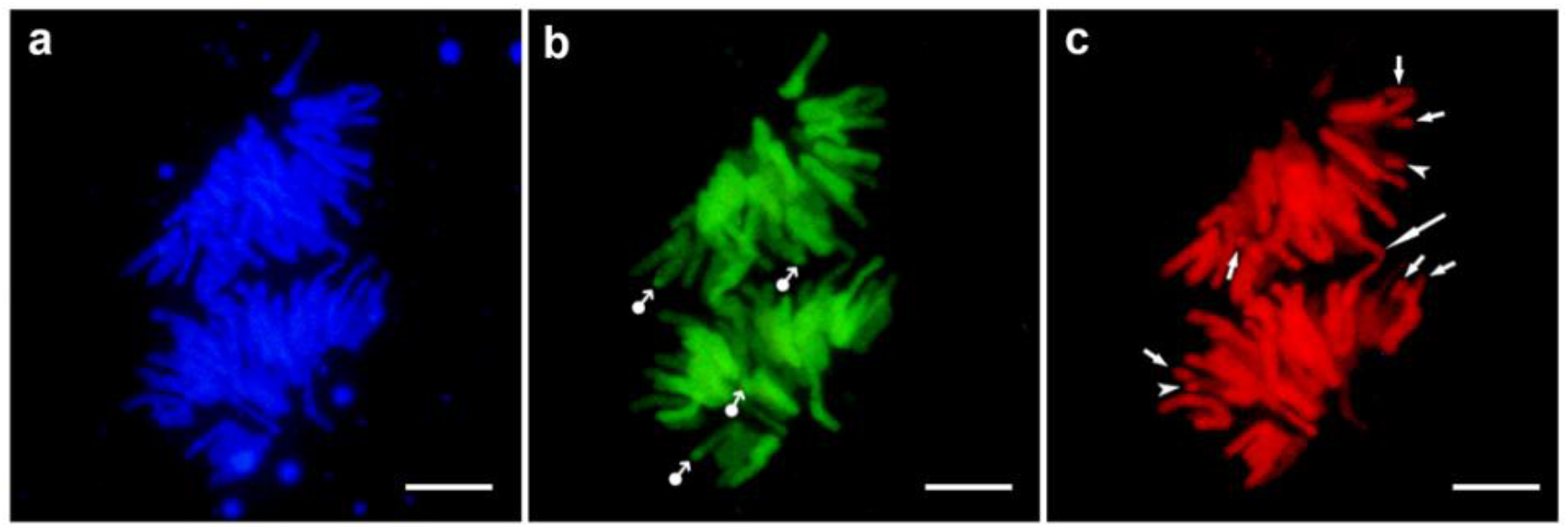
Chromosome rearrangements in Avena maroccana detected by GISH. Pictures are related to Fig. 2c in the article, showing studied anaphase in three different filters. a – DAPI-stained chromosomes; b – FITC-detected Avena nuda probe (As genome); c – TRITC-detected Avena eriantha probe (Cp genome). Translocations are marked by dot-arrows for terminal A/C, arrows for terminal C/A, and arrowheads for subterminal C/A. The large arrow indicates the bridge. Scale bars 10 µm

